# Three-dimensional chromatin organisation shapes origin activation and replication fork directionality

**DOI:** 10.1101/2022.06.24.497492

**Authors:** Katherine A. Giles, Noa Lamm, Phillippa C. Taberlay, Anthony J. Cesare

## Abstract

Faithful DNA replication requires the orderly firing of replication origins across the genome. At present, we lack details around how origins are selected for activation and the subsequent impact of this on replication dynamics. Here, we have investigated how chromatin organisation contributes to replication initiation and dynamics by intersecting ChIP-seq, Hi-C, Repli-seq, and OK-seq data from primary and tumour cells lines. We found replication initiation is significantly enriched at TAD boundaries, co-localizing with CTCF and cohesin in early and mid S-phase. Strong replication fork directionality (RFD) from initiation zones in TAD boundaries could occur in a bi- or uni-directional manner, which highly correlated with replication timing. While TAD boundaries were largely invariant, a minority of initiation zones were shared across cell lines, indicative of cell type specific regulation. These data are consistent with chromatin structure organizing replication initiation and dynamics, ensuring orderly completion of replication from TAD boundaries into TAD internal regions.

## Introduction

Faithful DNA replication requires ordered copying of the genome once per cell cycle. During S-phase DNA replication initiates from specialised chromatin regions called origins (Jacob and Brenner, 1963). Prior to S, replication origin licencing commences in mitosis and continues through G1-phase. This process initiates with origin recognition complex (ORC) binding to the DNA, followed by CDC6 and CDT1 cofactor association with chromatin bound ORC, and culminating with recruitment of the minichromosome maintenance (MCM) replicative helicase complex to form the pre-replication complex (Ding and Koren, 2020). Once established, origins remain dormant until DNA synthesis initiates in S, during which a subset of licensed origins are activated in sequential fashion across the genome (Limas and Cook, 2019). The human genome contains between 30,000-50,000 licensed origins (Cvetic and Walter, 2005; Mechali, 2010) that are well spaced to ensure the entire genome is replicated. It remains unclear, however, how origins are recognised and selected for activation to ensure a timely S-phase (Marks et al., 2017).

Chromatin is a dynamic, three-dimensional (3D) structure comprised of DNA, histones, and other associated proteins. During interphase, chromatin is arranged into topological associated domains (TADs) that compartmentalise the genome into nuclear micro-environments. This facilitates DNA dependent functions, such as enhancer-promoter interactions for transcription (Achinger-Kawecka et al., 2020; Giles et al., 2019; Khoury et al., 2020; Taberlay et al., 2016; Wang et al., 2019; Zhang and Kutateladze, 2019). TADs also influence replication timing precision (Du et al., 2021; Pope et al., 2014). This is regulated in part through TAD organization in 3D space, with early replicating TADs positioned at the nuclear centre and late replicating TADs at the nuclear periphery (Dimitrova and Gilbert, 1999). While the replication program is highly linked to 3D chromatin organisation, local chromatin features also exert influence on replication dynamics. For example, early replicating TADs contain cis-acting elements with enhancer like properties that govern domain-wide control over replication timing (Sima et al., 2019). The intimate connection between chromatin organisation and replication timing establishes a functional role, yet it remains unclear how 3D chromatin structure is linked to initiation events or replication fork dynamics during DNA synthesis.

Here, we have identified connectivity between 3D chromatin organisation, origin activation, replication fork directionality, and replication timing in both cancer and primary human cells. We found that replication initiation is significantly enriched at TAD boundaries specifically in early and mid, but not late, S-phase. Examining temporal replication dynamics revealed a prominent correlation between replication fork directionality (RFD) and replication timing. Strong RFD correlated with decreased replication timing as a function of genetic distance from the initiation zone, indicative of persistent replication from the initiation site into the TAD as S-phase progresses. Conversely weak RFD correlated with minimal change in replication timing indicative of local forks moving in both directions for shorter distances. Strong RFD scores could be bi- or uni-directional from an individual initiation zone. Cohesin, CTCF, and chromatin accessible sites were significantly enriched at early and mid-initiation zones, regardless of strong or weak RFD, but were not enriched at late initiation zones. While HeLa carcinoma, K562 leukemia lymphoblast, and IMR90 primary fibroblasts displayed common TAD boundary initiation zones, most initiation sites were not shared across all three cell types. Together these data expand the prior observation that replication initiates at TAD boundaries (Emerson et al., 2022; Lombardi and Tarsounas, 2020; Petryk et al., 2016) and reveal a broad functional role for 3D chromatin organisation in replication dynamics with cell type specific regulation.

## Results

### Early- and mid-S replication initiation zones are associated with TAD boundaries

Local chromatin organisation contributes to the spatiotemporal replication program (Audit et al., 2009; Guillou et al., 2010; Smith and Aladjem, 2014; Su et al., 2020). In concordance, we found that the architectural chromatin factors cohesin (Rad21 and SMC3 subunits) and CTCF, as well as chromatin accessible sites, were highly colocalised with ORC binding (Figure 1A-B) in publicly available ChIP-seq and DNase-seq data sets from HeLa cervical carcinoma and K562 leukemia lymphoblast cells (Supplementary Table 1). Cohesin and CTCF have diverse chromatin architectural roles beyond local chromatin organisation, including TAD formation (Nuebler et al., 2018; Rao et al., 2014; Wutz et al., 2017), and we thus investigated possible connections between replication origins and 3D chromatin architecture.

**Figure 1.**
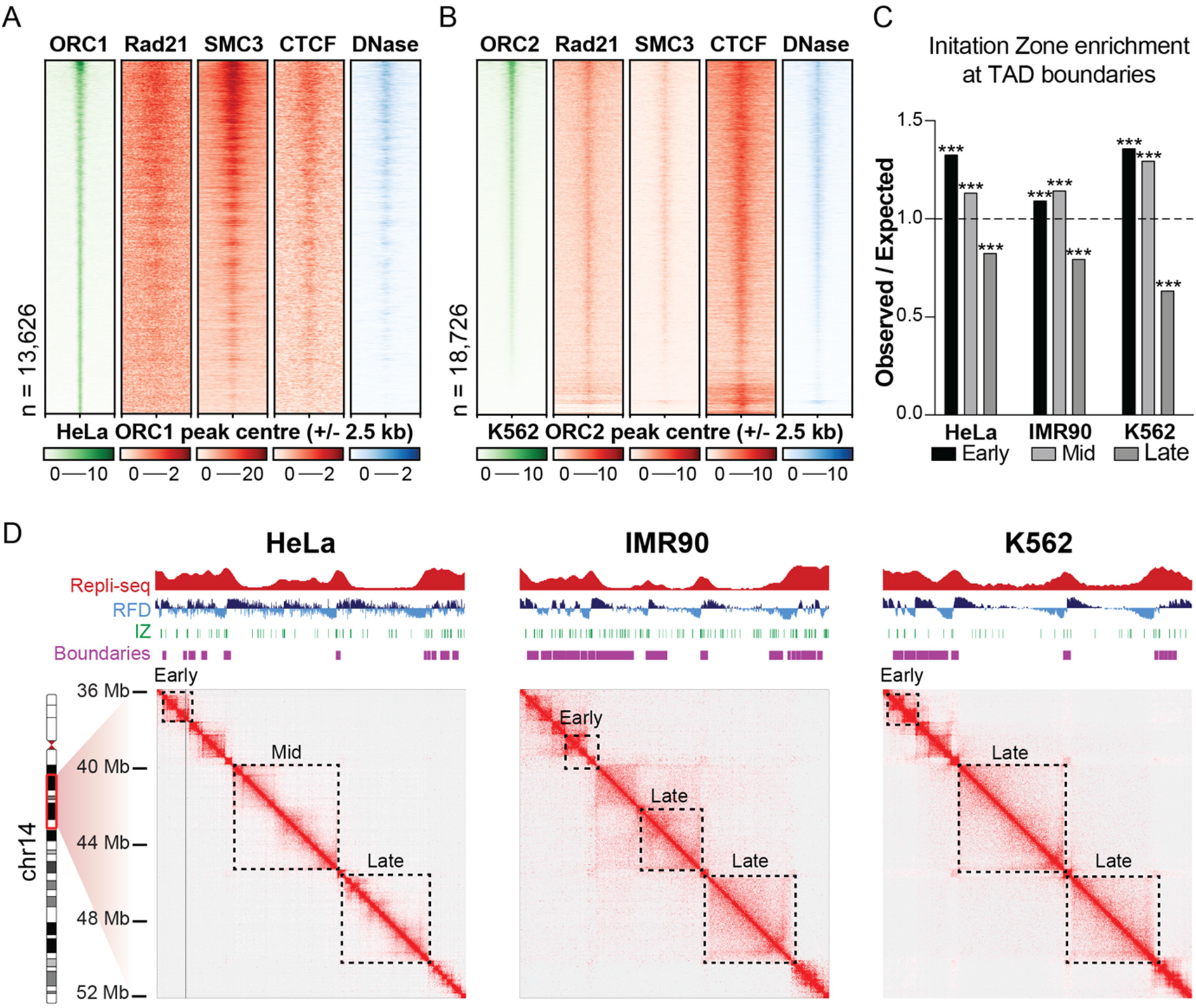
Early and mid-S replication initiation zones are associated with TAD boundaries. **(A)** Heatmap of ORC1 binding sites in HeLa cells, with corresponding ChIP-seq signal for Rad21, SMC3 and CTCF, and DNase accessible regions. Regions are ordered according to ORC1 signal strength and plotted +/- 2.5 kb from the centre of the ORC1 binding site. **(B)** As in (A) for ORC2 binding sites in K562 cells. **(C)** Observed / expected value for initiation zones in early, mid and late replicating regions at TAD boundaries. Initiation zones are significantly enriched with a score above one or significantly depleted with a score below one. A Benjamini–Hochberg adj p-value of *p<0*.*0001* is denoted as ***. **(D)** Example region of replication timing (RepliSeq; score scale 0-100), replication fork directionality (RFD; score scale −1 to 1), initiation zones, TAD boundaries and Hi-C contact maps on chromosome 14 from 36-52 Mb. Dotted boxes outline early, mid and late replicating TADs.

To do so we compared publicly available Hi-C, Repli-seq, and OK-seq data derived from HeLa, K562, and IMR90 primary fibroblast cultures (Supplementary Table 1). Hi-C identifies 3D chromatin organization including TAD architecture, Repli-seq reveals when during S-phase segments of DNA are replicated (replication timing), and OK-seq identifies the location of newly synthesised Okazaki fragments on the discontinuous lagging-strand during DNA replication (Burgers and Kunkel, 2017; Liu et al., 2022; Petryk et al., 2016). Mapping Okazaki fragments from OK-seq in a strand specific manner identifies replication fork direction, which is used to delineate replication initiation and termination zones (Liu et al., 2022; Petryk et al., 2016). Initiation zones are regions of active replication initiation, in contrast to ORC binding, which delineates both active origins and those which are licenced but remain dormant.

Intersecting replication initiation zones, TAD architecture, and replication timing, revealed a strong association between these features. In all three cell lines examined replication initiation was significantly enriched (observed/expected > 1; Benjamini–Hochberg adj p-value *p<0*.*0001*) at TAD boundaries, which was strongly influenced by early and mid S-phase progression (Figure 1C). Specifically, 71-90% of replication initiation zones in early S (HeLa 71%, IMR90 90% and K562 79%), and 58-78% of replication initiation zones in mid S (HeLa 58%, IMR90 78% and K562 68%), were located within TAD boundaries (Supplementary Figure 1A-C). Conversely, in late S-phase 54-72% of replication initiation zones were internalized within TADs (HeLa 57%, IMR90 54% and K562 72%; Supplementary Figure 1A-C) and were significantly depleted from TAD boundaries (observed/expected < 1; Benjamini– Hochberg adj p-value *p<0*.*0001*; Figure 1C). Replication thus initiates most often in early and mid-S from TAD boundaries, and as S-phase progresses the diminishing number of initiation events occur with a greater propensity inside TADs domains. Additionally, we observed TADs with late replicating internal regions whose boundaries exhibited early- or mid-S initiation events (as illustrated on chromosome 14, Figure 1D), consistent with replication progression from the boundary into the TAD as S-phase continues.

### Replication fork directionality and timing correlate with initiation events occurring at TAD boundaries

To gain a better understanding of TAD replication dynamics, we examined replication fork directionality and timing in the 100 kb on either side of replication initiation zones within TAD boundaries (Figure 2A-C and Supplementary Figures 2A-C, 3A-C). We defined strong RFD as a mean score of < −0.2 (left moving) or > +0.2 (right moving), and weak RFD as scores between −0.2 and +0.2. Intersected with RFD were Repli-seq data revealing when in S-phase the DNA sequences in the 100 kb region flanking regions were replicated.

**Figure 2.**
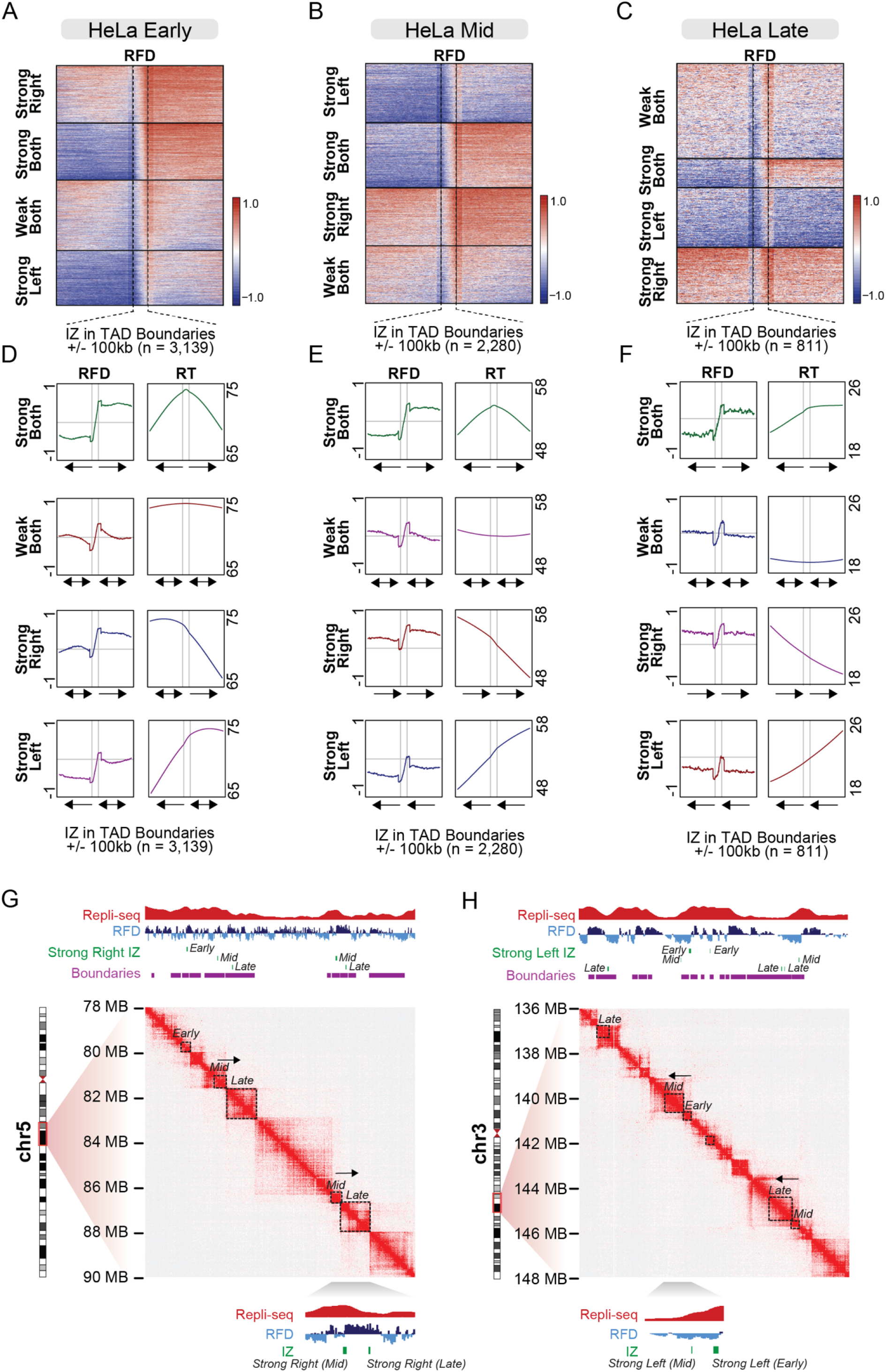
Replication fork directionality initiating from TAD boundaries. **(A-C)** Heatmaps of RFD in early (A) mid (B) or late (C) replicating regions which overlap with TAD boundaries. Initiation zones are scaled to line up the boarders of each region. RFD is plotted +/- 100 kb either side of the initiation zone, clustered in four groups by kmeans clustering, with each cluster sorted in descending signal order. **(D-F)** Profile plots of RFD and replication timing (RT) within each of the four clusters in (A-C). Arrows indicate average RFD either side of the initiation zone. Vertical grey lines indicate initiation zone boarders and horizontal grey line indicates a score of zero for RFD. **(G, H)** Example region of replication timing (Repli-seq; score scale 0-100), replication fork directionality (RFD; score scale −1 to 1), initiation zones of strong right moving forks (G) or strong left moving forks (H), TAD boundaries and Hi-C contact maps on chromosome 5 from 78-90 Mb (G) and chromosome 3 136-148 Mb (H). Dotted boxes outline early, mid and late replicating TADs where there is strong RFD from a mid to late replicating TAD (G) and early to mid-replicating TAD (H). An example of each is expanded below each contact map. Also see Supplementary Figures 2 and 3.

In all three cell lines we identified a subset of TAD boundaries with DNA replication initiation in early, mid, and late S-phase that presented strong RFD scores (Figure 2A-C and Supplementary Figures 2A-C, 3A-C). These strong RFD signals were accompanied by a progressive reduction in replication timing as a function of genetic distance from the initiation zone (Figure 2D-F and Supplementary Figures 2D-F, 3D-F). Conversely, when early- and mid-S initiation zones within TAD boundaries displayed weak RFD scores, this correlated to a minimal decrease in replication timing score as a function of distance from the initiation zone (Figure 2D-F). Weak or strong RFD from early- and mid-S phase initiation zones at TAD boundaries occurred in both a bi- and uni-directional manner, resulting in four classifications (strong both, strong right, strong left, weak both).

The implication of these data are that the strength of replication fork directionality from initiation events within TAD boundaries directly correlates with the propensity for replication forks to advance a long-distance into the adjacent TAD as S-phase progresses. This outcome results in the progressive diminishment of replication timing moving away from the initiating zone as replication persists through S-phase. Contrarily, weak RFD from origins in TAD boundaries predominantly corresponded with minimal reduction in replication timing as a function of genetic distance from the initiation zone. The implication of this observation is that initiation zones with weak RFD have nearby forks concomitantly moving towards the weak initiation zone that diminish RFD score, result in timely completion of replication in the nearby genomic regions, and produce more frequent termination events. Examples of weak RFD from TAD boundaries are demonstrated in the early replicating TADs on chromosome 14 in Figure 1D, while strong RFD is demonstrated from the boundaries of the late replicating TADs. Initiation zones displaying strong uni-directional RFD suggest the strength of replication directionality and persistence is not an intrinsic property of an individual origin, but instead impacted by chromatin features in the adjacent region.

The characteristics of strong uni-directional RFD from initiation zones within TAD boundaries varied dependent on S-phase stage. In mid and late S we observed initiation zones with strong right moving uni-directional RFD, adjacent to a downstream initiation zone on the right, which also exhibited a strong uni-directional rightward RFD. Similar events were observed with adjacent strong left moving uni-directional RFD (Figure 2B, 2E and Supplementary Figures 2B, 2E, 3B, 3E). We sought to identify if adjacent initiation zones with strong uni-directional RFD in the same direction exhibited concurrent or progressive replication timing. We indeed found examples, such as on chromosome 5 (Figure 2G), where a mid-replicating boundary with a strong right moving fork had a mid-replicating TAD on the left and a late replicating TAD on the right. Similar was identified for strong left moving forks on chromosome 3, between early to mid replicating TADs and as mid to late TADs (Figure 2H). In such events, the juxtaposition of forks with strong uni-directional RFD moving in the same orientation, with progressive reduced replication timing scores, is consistent with replication persisting with stable directional progression from initiation at the TAD boundary and into the TAD domain as S-phase progresses. Similar events were not common during early S-phase.

Similarity of outcomes across HeLa, K562 and IMR90 cells (Figure 2, and Supplementary Figures 2 and 3) indicates conservation of RFD and replication timing in diverse cells, consistent with 3D chromatin structure regulating replication outcomes.

### Chromatin architecture proteins are enriched at all initiation zones regardless of RFD

We also examined the enrichment of ORC, CTCF, cohesin subunits (Rad21 and SMC3), and chromatin accessible sites, at initiation zones in HeLa and K562 cells. We find that early and mid S-phase initiation zones were significantly enriched for all these features regardless of strong or weak RFD in both cell lines (Supplementary Figure 4A-D). Late S-phase initiation zones in K562, however, displayed significant enrichment of ORC2 and DNase sites, but cohesin and CTCF were neither significantly enriched nor depleted (Supplementary Figure 4E). Conversely, in HeLa cells, ORC, cohesin, CTCF and DNase sites were all significantly depleted from strong uni-directional or weak bidirectional late S-phase initiation zones (Supplementary Figure 4F). Together these data suggests that local chromatin structure at active origins is linked to replication timing. Early and mid-replicating initiation zones are enriched for chromatin architectural proteins in both cell lines examined, while late replicating regions display cell type differences and may be void of these chromatin features. Further, the presence or absence of these features are independent of the role of 3D compartmentalising the genome for replication fork directionality, as we find no difference in their enrichment for each RFD initiation group in early and mid S-phase.

### Replication initiation zones are both common and cell type specific

Given that TADs are largely invariant across different cell types (McArthur and Capra, 2021; Smith et al., 2016a), we next sought to determine the commonality of chromatin architecture and replication dynamics across HeLa, IMR90 and K562 and identify if this has an impact on replication initiation. We first intersected TAD boundaries and found 87% of the HeLa boundaries overlapped with 50% of the IMR90 boundaries and 65% of the K562 boundaries (Figure 3A). We performed the same analysis on initiation zones and found that the percentage of overlap of initiation zones was much lower; 32% of HeLa initiation zones overlapped with 25% of IMR90 and 44% of K562 (Supplementary Figure 5A). This overlap was even lower for initiation zones occurring within TAD boundaries (Hela 25%, IMR90 18% and K562 33%; Figure 3B). Therefore, while all cell lines display similar enrichment of replication initiation from TAD boundaries, the genomic position of the initiation zone within the boundary is variable between cell types.

**Figure 3.**
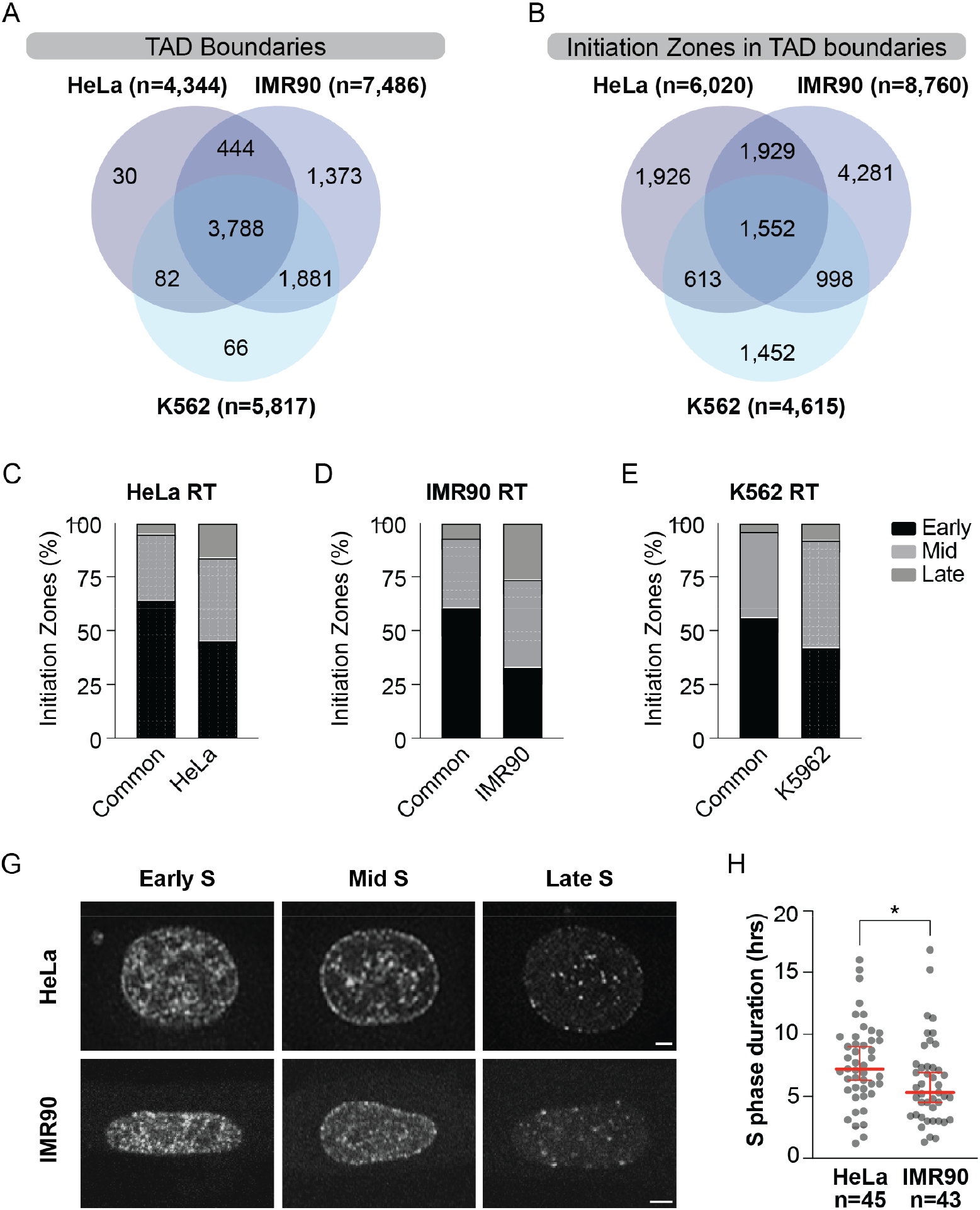
Cells maintain similar spatiotemporal replication programs despite cell type specific differences to initiation zones. **(A)** Venn diagram of TAD boundary intersections between HeLa, IMR90 and K562 cells. **(B)** Venn diagram overlap of initiation zones within TAD boundaries for HeLa, IMR90 and K562. **(C, D, E)** Initiation zones within TAD boundaries for those common to HeLa, IMR90 and K562 and initiation zones that are in each cell line, but not common to all cell lines. Bars show the percentage of initiation zones in early, mid and late S-phase for the replication timing (RT) profile for the respective cell line. **(G)** Representative images of PCNA foci from early, mid and late S-phase cells in HeLa and IMR90s. Scale bar = 5µM. **(H)** S-phase duration (hours) for HeLa (n=45) and IMR90 (n=43) cells. Red line represents mean and error bars are SD. Significance is \**p=0*.*0169*, two-tailed Mann Whitney non-parametric T-test.

We next focused on how the common and cell type specific initiation zones at TAD boundaries were activated across S-phase for each cell type. We found that of the initiation zones common to all cell lines ∼ 60% were activated in early S-phase and less than 6% were activated in late S-phase (Figure 3C-E). The remainder of the initiation zones that were either cell type specific, or present in two of the three cell lines, had a higher percentage of activation in late S-phase (7-26%) as compared to the common initiation zones (Figure 3C-E). This suggests that the common initiation zones in TAD boundaries are important for early replication events, whereas cell-type specific initiation zones become more prominent in the later stages of replication.

We also analysed S-phase length and the spatiotemporal pattern of replication by tracking PCNA foci in HeLa and IMR90 cultures using live cell imaging. Despite differences in replication initiation observed in the sequencing data, the spatiotemporal pattern of replication was similar between both cell types (Figure 3G). There was, however, a significant difference in S-phase length, which was shorter in IMR90 as compared to HeLa (Figure 3H). Of note IMR90s had an increased number of TAD boundaries and initiation zones compared to HeLa cells consistent with a greater number of fired origins per S-phase.

## Discussion

DNA replication is regulated on multiple levels to ensure accurate copying of the DNA once per cell cycle. Here we expand on the role of 3D chromatin organisation in genome copying and show that chromatin structure impacts multiple facets of DNA replication. Our study demonstrates a link between active origin firing, replication dynamics, and 3D chromatin structure. We find DNA replication primarily initiates from TAD boundaries during early and mid S, but as the cell cycle proceeds into late S, replication initiation events within TAD domains become more prevalent. These outcomes were common in two cancer, and one primary cell type, indicative of conserved mechanisms in healthy and transformed cells. Notwithstanding conserved general regulatory mechanisms, we did observe cell-type specific outcomes. While the three cell lines analysed shared the majority of TAD domain structure, they possessed a minority of common initiation zones. Cell lines therefore display both common and specific regions of replication initiation (Smith and Aladjem, 2014; Smith et al., 2016b). Where initiation zones were common across all three cell types, these were most often active in early S, with only 6% of late S-phase initiation zones employed in HeLa, K652 and IMR90.

We also found a correlation between RFD and replication timing. Strong RFD correlated with persistent decreasing replication fork timing as forks moved away from the initiation zone. This is consistent with long range replication events as cells progress through S-phase. Weak RFD correlated with minor changes in replication timing. These data are consistent with short range replication events, on-coming forks in the other direction completing replication in the adjacent region, and more common replication termination. Strong or weak RFD could be bi- or uni-directional from an individual initiation zone, indicating the quality determining RFD strength was not the initiation zone per se, but some other chromatin quality in the surrounding region.

We observed ORC, CTCF and cohesin binding, along with DNase accessible regions, were significantly enriched in early and mid-replicating boundaries in both HeLa and K562. In late S-phase, however, these features were significantly depleted from HeLa initiation zones, and neither enriched nor depleted in K562. Notably, the presence or absence of CTCF, cohesin, or DNase accessible regions had no impact on the strength of RFD. While these data suggest none of these factors influence RFD strength, we cannot rule out a nuanced role for these features in controlling replication dynamics. Possibilities include CTCF directionality or nucleosome phasing exerting influence over origin firing and RFD strength. For example, high-density arrays of CTCF motifs were recently identified at high-efficiency early S-phase initiation zones (Emerson et al., 2022). CTCF deletion, however, did not significantly alter replication timing (Sima et al., 2019), suggesting CTCF is likely not the driving factor of origin firing efficiency. Further research into the role of CTCF and other architectural proteins is needed to fully understand their potential roles regulating DNA replication through chromatin organisation.

There is an increasing body of data showing a role for epigenetics in both DNA replication and 3D chromatin structure. TAD boundaries and ORC binding are enriched for active histone marks such as H3K4me3 and H3K27ac (Pelham-Webb et al., 2021; Sima et al., 2019; Taberlay et al., 2016). Additionally, early replicating control elements are enriched for enhancer like epigenetic marks (Sima et al., 2019). More recently, loss of DNA methylation was associated with altered precision of replication timing and 3D chromatin structure (Du et al., 2021). These links highlight the combined influence of the epigenome and 3D chromatin structure on replication timing. However, whether the epigenome has a role specifically in priming chromatin for replication and origin licencing, or if it additionally functions in selecting origins for activation is not known. It is interesting to speculate that epigenetic control of these processes is what allows flexibility in the replication program of different cell types or the specific bi- and uni-directional RFD strength from initiation zones reported above.

A major and intriguing finding of this study was our identification of arrayed initiation zones within TAD boundaries with common strong uni-directional rightward or leftward RFD. These regions were linked to a replication timing transition from early to mid, or mid to late, and consistent with a strong overall uni-directional replication. While the biochemical mechanism of semi-conservative DNA replication is bi-directional, this does not demand replication proceed equidistant in both directions within the chromatin context. We predict as origins and TADs are established in mitosis and G1, TAD boundaries are selected as the prominent location for replication initiation early in the oncoming S-phase. These early replication regions may be common or specific between cell lines, depending on the similarity or dissimilarity of chromatin architecture. Local chromatin features dictate RFD strength from the early and mid-S TAD boundary initiation zones, with some regions having more prominent left or rightward moving forks. In some regions, RFD is arrayed to ensure progressive completion of early and mid-replicating regions before late origin firing occurs. Because replication initiates primarily from TAD boundaries in early and mid-S, the final regions of the genome copied are internal within TADs. Which late-S initiation zones withing TADs are utilised is likely influenced in a stochastic nature depending on how replication has progressed thus far in S-phase. We expect this stochastic nature is why late-S initiation zones are not commonly shared and display a lower overall lower RFD.

In summary, our study links replication dynamics with 3D chromatin organisation. Our findings support TAD boundaries as having a significant role for replication initiation in early and mid S-phase and establish a role for 3D chromatin organisation in replication fork directionality. The consistency of our findings across both cancer and primary cells, highlight the conservation of this essential function. Future efforts are required to understand how the cell regulates specific RFD strength at individual initiations zones. Further, how 3D chromatin organization impacts replication dynamics during oncogenic or chemotherapeutic induced replication stress is of interest.

## Acknowledgements

We thank Will Hughes and the Advanced Microscopy Centre at the Children’s Medical Research Institute for providing the infrastructure and technical support for microscopy experiments. P.C.T. is supported by Australian National Health & Medical Research Council (NHMRC) Investigator (GNT1176417), and Project (GNT1161985) grants. A.J.C is supported by an Australian Research Council (ARC) Future Fellowship (FT210100858), and work in his lab is supported by grants from the ARC (DP210103885) and NHMRC (1185870).

## Author Contributions

Conceptualization – K.A.G., P.C.T and A.J.C.

Formal Analysis – K.A.G.

Investigation – K.A.G. and N.L.

Writing – Original Draft – K.A.G

Writing – Review & Editing – K.A.G., N.L., P.C.T., A.J.C.

Visualization – K.A.G., N.L.

Supervision – P.C.T. and A.J.C.

Funding Acquisition – P.C.T. and A.J.C.

## Inclusion and diversity

We worked to ensure diversity in experimental samples through the selection of cell lines and genomic data sets. One or more of the authors of this paper received support from a program designed to increase minority representation in science.

## STAR Methods

| REAGENT or RESOURCE | SOURCE | IDENTIFIER |
| --- | --- | --- |
| **Laboratory consumables** |  |  |
| Prolong gold antifade mounting media | ThermoFisher Scientific | P36930 |
| Immersion oil | Zeiss | 444970-9010-000 |
| DMEM culture medium | Gibco | 11965-092 |
| FluoroBrite DMEM | Gibco | A18967-01 |
| GlutaMax | Gibco | 35050-061 |
| Non-essential amino acids | Gibco | 11140-050 |
| Bovine growth serum | Hyclone | SH30541.03 |
| Fetal bovine serum | Sigma | F9243 |
| MycoAlet | Lonza | LT07-118 |
| Tissue Culture Dish with Cover Glass bottom | World Precision Instruments | 22092019 |
| Cell Cycle RFP-PCNA chroobody | Chromotek | CCR |
| Lipofectamine LTX | Thermo Fisher Scientific | 15338-100 |
| **Laboratory Equipment** |  |  |
| Zeiss Cell Observer SD spinning-disk confocal microscope | Zeiss |  |

| **Software** |  |
| --- | --- |
| Software / algorithm name | Source |
| ZEISS Zen Black and Zen Blue | Zeiss (v2.0.14283.302) |
| Graphpad Prism v9 | GraphPad Software (v9.4.0) |
| Adobe Illustrator 2022 | Adobe (v26.3.1) |
| Adobe Photoshop 2022 | Adobe (v23.3.2) |
| R | The R Foundation (v3.6.2) |
| Seqplots | http://przemol.github.io/seqplots/# (v1.10.6) |
| JuiceBox | https://aidenlab.org/juicebox/ (v2.1.10) |
| IGV | Broad Institute (v2.8.0) |

## Resource availability

Further information and requests for resources should be directed to the Lead Contact.

### Materials availability

This study did not generate new unique reagents.

## Method details

### Data and Code Availability

All next generation sequencing datasets used in this study have been previously published (Dellino et al., 2013; Miotto et al., 2016; Petryk et al., 2016; Pope et al., 2014; Rao et al., 2014). Raw and processed files for ChIP-seq, DNase-seq, Hi-C and Repli-seq are available on Gene Expression Omnibus (ncbi.nlm.nih.gov/geo/), with accession numbers listed in Supplementary Table 1. Processed OK-seq data (RFD, initiation zones and termination zones) is available on Github (github.com/CL-CHEN-Lab/OK-Seq). All code used for analysis and data visualisation is available from public sources and packages available on Bioconductor and Github.

### Data Intersections and enrichment

Data intersections were performed with R with *GenomicRanges* package v1.38.0. Significant enrichment or depletion was determined with the genome association tester v1.0 (Heger et al., 2013). The observed over expected value was calculated with 10,000 iterations and determine statistically significant if the q-value was less than 0.0001.

### Data visualisation

ChIP-seq and DNase-seq heatmaps were generated with deepTools2 v3.5.1. RFD heatmaps and profile plots were generated with Seqplots v1.10.6. Hi-C contact maps were created in JuiceBox v2.1.10. Genome track images of OK-seq, Repli-seq, domain boundaries and initiation zones were generated with IGV v2.8.0.

### Cell Culture Maintenance

HeLa cells and IMR90 fibroblasts were kindly provide by J. Karlseder (Salk Institute). All cells were grown in DMEM (Gibco, 11965-092) supplemented with and 1% GlutaMax (Gibco, 35050-061) and 1% non-essential amino acids (Gibco, 11140-050). HeLa cell culture media was supplemented with 10% bovine growth serum (Hyclone, SH30541.03) and IMR90 culture media with 10% fetal bovine serum (Sigma, F9243). Cell cultures were maintained in 37 ^º^C with 10% CO_2_ and 3% O_2_. Cells were passaged at approximately 80% confluence. Both cell lines were verified by Cell Bank Australia using STR profiling and confirmed to be mycoplasma negative (MycoAlet, LT07-118, Lonza).

### Live Cell Imaging

Cells were prepared for live cell imaging by seeding 2 × 10^5^ cells on a 3 cm glass bottom culture dish (World Precision Instruments). After 24 hrs cells were transfected with RFP-PCNA chromobody (Chromotek, PCNA-CB, purchased with a material transfer agreement) using Lipofectamine LTX (Thermo Fisher Scientific, 15338-100) as per the manufacturer’s instructions. After 24 hrs media was replaced with FluoroBrite DMEM (Gibco, A18967-01), supplemented as described above. Imaging was performed in culture conditions described above on a Zeiss Cell Observer SD spinning-disk confocal microscope using combined differential interface contrast and fluorescence imaging (a 561 laser, 7% excitation power, 1 × 1 binning, and EM gain of 920) with appropriate filter sets and a x63 / 1.3 NA oil-immersion objective. Ten images per z stack (9 *μ*m) were captured in an image scaled to 47,504 px x 37,602px at 10.05 µM using Zen Blue 2 (Zeiss, v2.0.14283.302) and Evolve Delta (Photometrics) camera every 90-200 seconds for up to 48 hrs. S-phase duration was calculated as time in hours from the appearance of the first PCNA foci until the disappearance of the last foci.

## Supplemental Figures and Tables

**Supplementary Figure 1.**
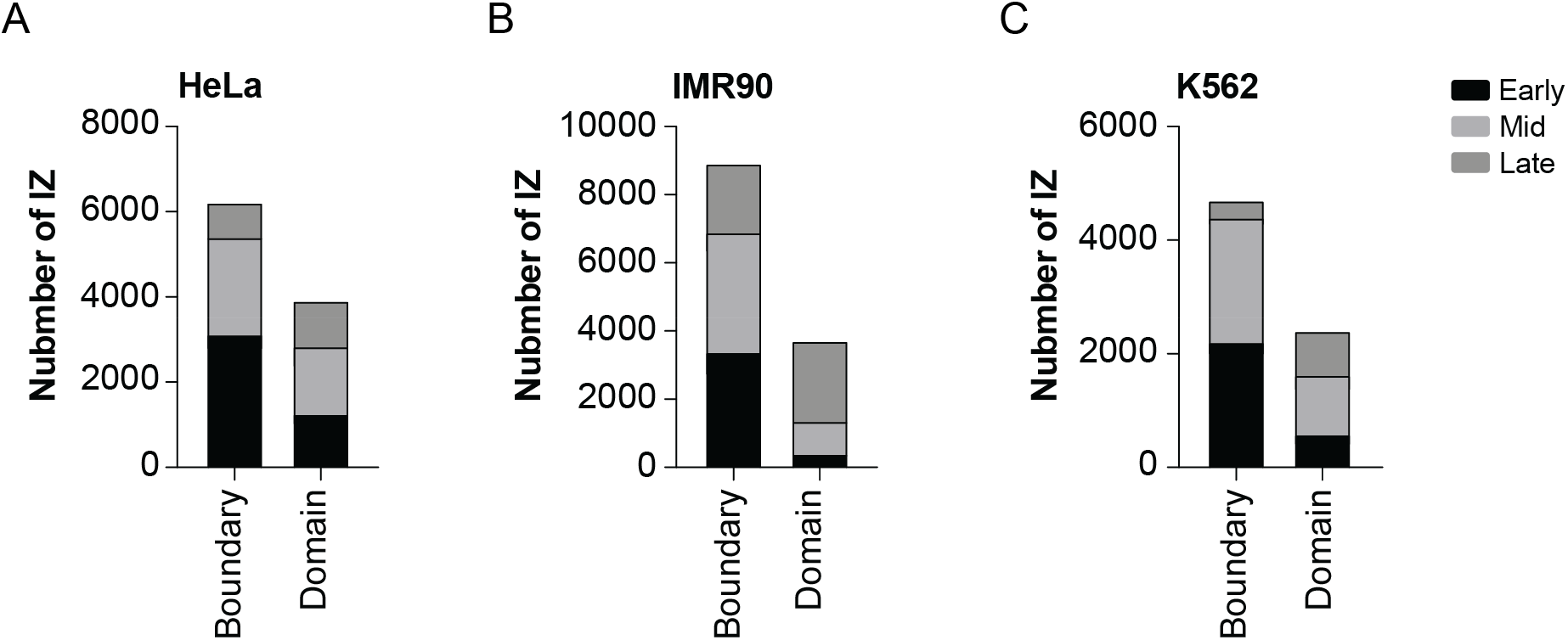
**(A)** Number of HeLa initiation zones overlapping with domain boundaries and TAD domains from HeLa Hi-C data. Initiation zones are subdivided into those overlapping early, mid or late replicating regions from HeLa Repliseq data. **(B)** As in (A) for IMR90. **(C)** as in (A) for K562.

**Supplementary Figure 2.**
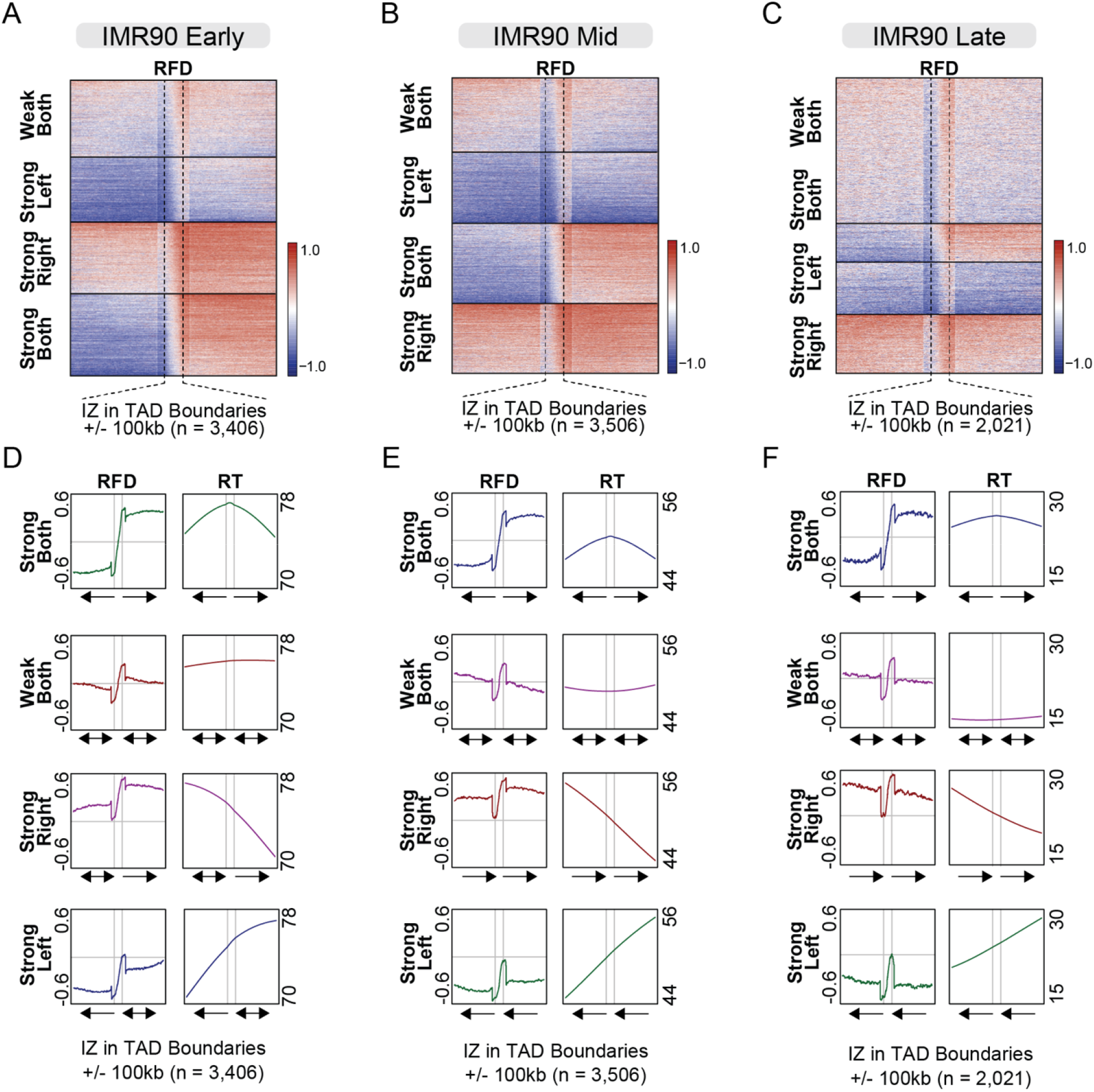
**(A-C)** Heatmaps of IMR90 replication fork directionality (RFD) in early (A) mid (B) or late (C) replicating initiation zones that overlap with TAD boundaries. Initiation zones are scaled to line up the boarders so that RFD either side of the initiation zone is plotted +/- 100 kb. Vertical dotted lines indicate initiation zone boarder. Initiation zones are clustered in four groups by kmeans clustering, with each cluster sorted in descending signal order. **(D-F)** Profile plots of IMR90 replication fork directionality (RFD) and replicating timing (RT) within each of the four clusters (from A-C). Grey lines indicate initiation zone boarders, with arrows indicating main RFD either side of the initiation zone. Horizontal grey line indicates a score of zero for RFD. Arrows under plots indicate average RFD.

**Supplementary Figure 3.**
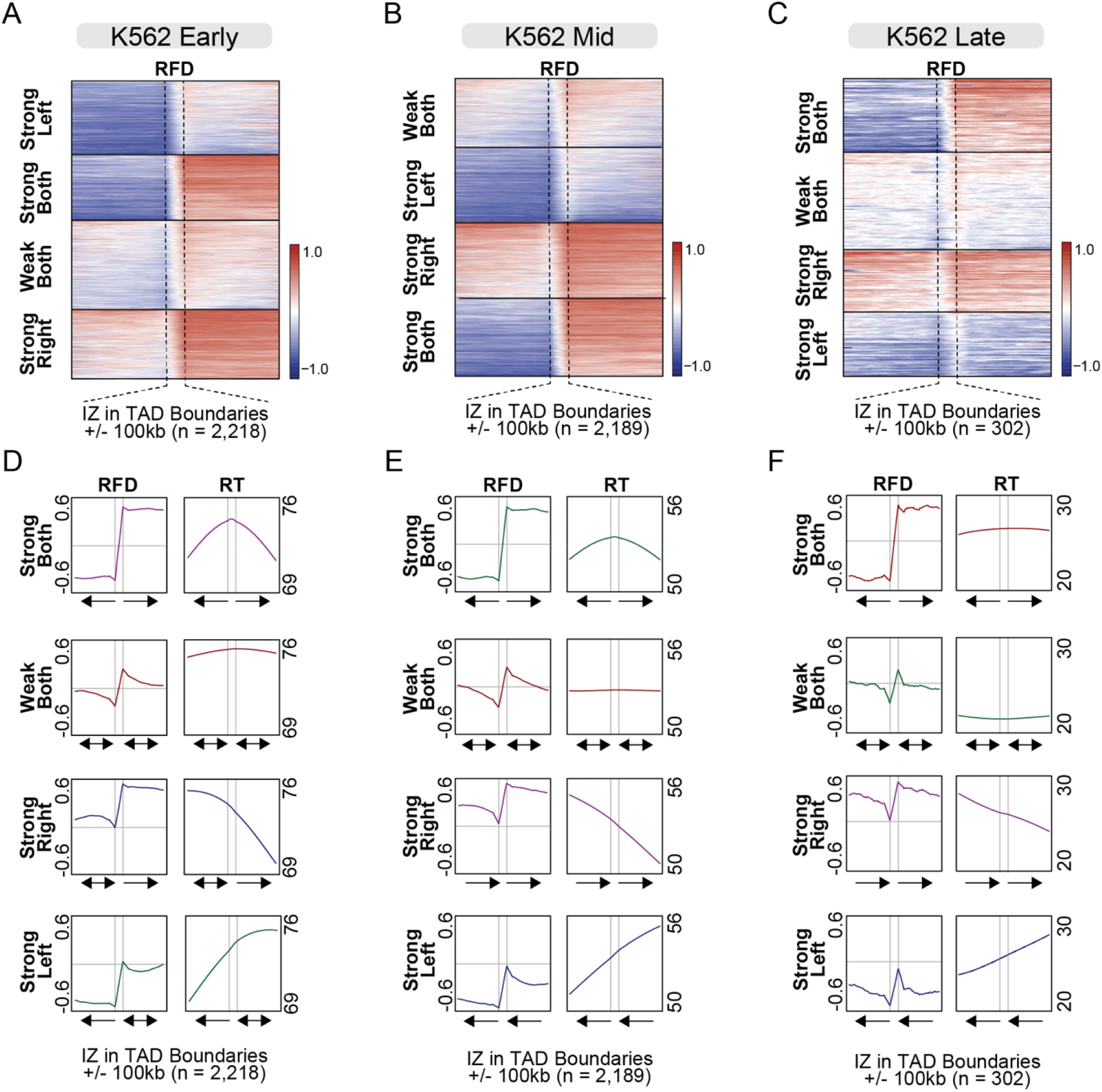
**(A-C)** Heatmaps of K562 replication fork directionality (RFD) in early (A) mid (B) or late (C) replicating initiation zones that overlap with TAD boundaries. Initiation zones are scaled to line up the boarders so that RFD either side of the initiation zone is plotted +/- 100 kb. Vertical dotted lines indicate initiation zone boarder. Initiation zones are clustered in four groups by kmeans clustering, with each cluster sorted in descending signal order. **(D-F)** Profile plots of K562 replication fork directionality (RFD) and replicating timing (RT) within each of the four clusters (from A-C). Grey lines indicate initiation zone boarders, with arrows indicating main RFD either side of the initiation zone. Horizontal grey line indicates a score of zero for RFD. Arrows under plots indicate average RFD.

**Supplementary Figure 4.**
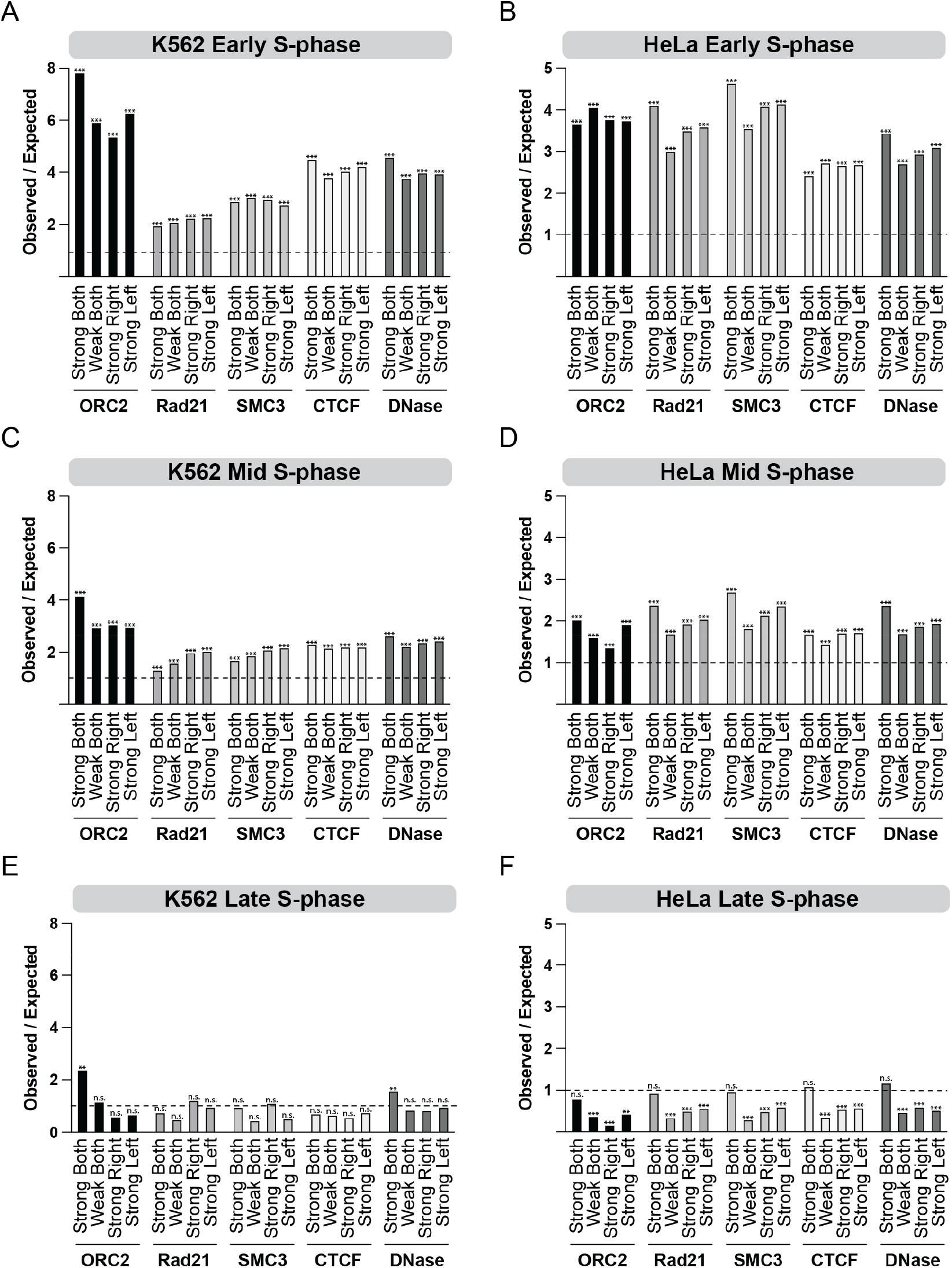
**(A-F)** Observed / expected enrichment of ORC2, Rad21, SMC3, CTCF and DNase accessible chromatin regions at initiation zones within TAD boundaries in K562 (A, C, E) and HeLa (B, D, F) cells. Initiation zones are subdivided into early (A-B), mid (C-D) and late (E-F) S-phase, and further subdivided based on replication fork direction from the groups in Supplementary Figure 3 (K562) and Figure 2 (HeLa). Initiation zones are significantly enriched with a score above one or significantly depleted with a score below one. A Benjamini–Hochberg adj p-value of *p<0*.*001* is denoted as **, *p<0*.*0001* as *** and not significant as ‘n.s.’.

**Supplementary Figure 5.**
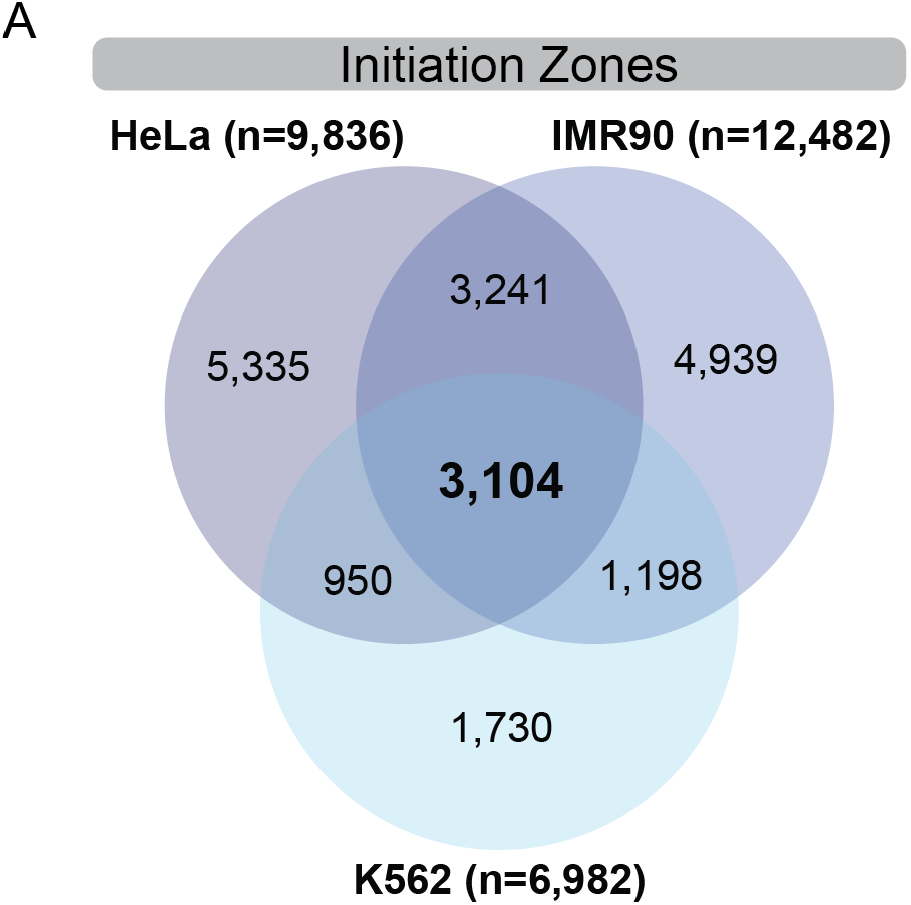
**(A)** Venn diagram of initiation zones intersections between HeLa, IMR90 and K562 cells.

**Supplementary Table 1.**
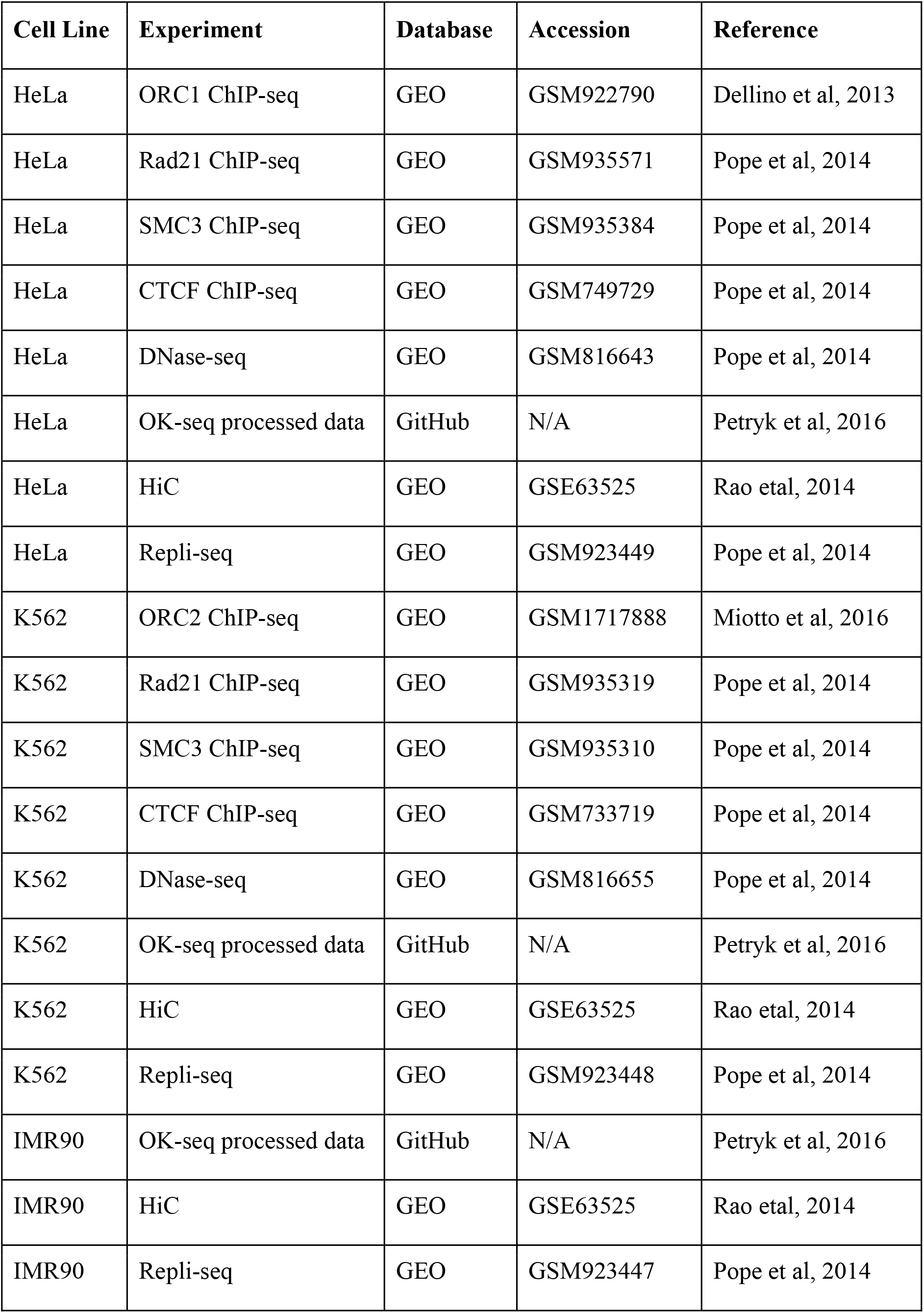
Accession numbers for published data used in this manuscript.

